# The scaling of genome size and cell size limits maximum rates of photosynthesis with implications for ecological strategies

**DOI:** 10.1101/619585

**Authors:** Adam B. Roddy, Guillaume Théroux-Rancourt, Tito Abbo, Joseph W. Benedetti, Craig R. Brodersen, Mariana Castro, Silvia Castro, Austin B. Gilbride, Brook Jensen, Guo-Feng Jiang, John A. Perkins, Sally D. Perkins, João Loureiro, Zuhah Syed, R. Alexander Thompson, Sara E. Kuebbing, Kevin A. Simonin

## Abstract

A central challenge in plant ecology is to define the major axes of plant functional variation with direct consequences for fitness. Central to the three main components of plant fitness (growth, survival, and reproduction) is the rate of metabolic conversion of CO_2_ into carbon that can be allocated to various structures and functions. Here we (1) argue that a primary constraint on the maximum rate of photosynthesis per unit leaf area is the size and packing density of cells and (2) show that variation in genome size is a strong predictor of cell sizes, packing densities, and the maximum rate of photosynthesis across terrestrial vascular plants. Regardless of the genic content associated with variation in genome size, the simple biophysical constraints of encapsulating the genome define the lower limit of cell size and the upper limit of cell packing densities, as well as the range of possible cell sizes and densities. Genome size, therefore, acts as a first-order constraint on carbon gain and is predicted to define the upper limits of allocation to growth, reproduction, and defense. The strong effects of genome size on metabolism, therefore, have broad implications for plant biogeography and for other theories of plant ecology, and suggest that selection on metabolism may have a role in genome size evolution.

## Introduction

Quantifying major axes of plant functional variation has given rise to an ever-growing list of traits that impact growth, reproduction, and survival, the three components of individual fitness (Violle et al. 2007). These traits have traditionally been viewed from a reductionist perspective that scales form-function relationships of individual plant organs (e.g. leaves, stems, and roots) to whole organism ecological strategies. As the ultimate source of energy and matter for growth and reproduction, photosynthetic capacity represents a first-order constraint on the emergent properties between whole plant form and function and individual fitness. Here we provide evidence that genome-cellular allometry directly influences interspecific variation in photosynthetic metabolism and provide a mechanistic framework that links genome size and metabolism to other aspects of plant ecology and evolution.

One of the three components of fitness is growth, which is ultimately limited by photosynthetic metabolism. Relative growth rate (RGR) varies considerably across species and is driven by photosynthetic rate and the resource investment to support photosynthesis as:

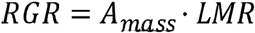

where *A*_*mass*_ is the photosynthetic rate per unit leaf biomass and LMR is the leaf mass ratio (the proportion of leaf dry mass to total plant dry mass). *A*_*mass*_ is, therefore, frequently considered a major plant strategy axis (Poorter and Remkes 1990; Poorter et al. 1990; Reich et al. 1992). However, *A*_*mass*_ can be decomposed as:

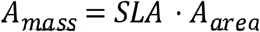

where SLA is the specific leaf area (leaf area per leaf dry mass) and *A*_*area*_ is the net carbon assimilation rate per unit canopy leaf area. Because of its direct effect on *A*_*mass*_, SLA is often considered a major predictor of interspecific variation in RGR. *A*_*area*_, on the other hand, varies orthogonally to SLA (Wright et al. 2004), and, therefore, determines the upper limit of the relationship between *A*_*mass*_, SLA, and RGR. Maximum potential *A*_*area*_ represents, then, a fundamental limitation on the maximum amount of carbon available for allocation to growth, reproduction, and survival relative to species ecological strategies.

The centrality of *A*_*area*_ to plant ecological strategy suggests two questions:

- First, what are the fundamental features of plant structure that determine maximum potential *A*_*area*_?
- Second, to what extent do these relationships scale up to affect plant ecological strategies and evolutionary dynamics?

Here we present a mechanistic framework to address both of these questions, that is based on the positive scaling between genome size and cell size. Although the relationship between genome size (i.e. nuclear volume) and cell size has long been of interest (von Sachs 1893), the mechanisms are still not fully understood (Doyle and Coate 2019), and its implications for organismal metabolism have not been fully articulated. We show that the allometry between genome size and cell size influences rates of photosynthetic metabolism and argue that the scaling of genome size and metabolism affects ecological distributions and evolutionary dynamics. In this way, any factor affecting rates of metabolism is a potential agent of selection on genome size and, potentially, on genome structure as well.

It is now widely recognized that variation in genome size can have significant consequences for organismal structure and function, independent of the genes that define the genotype (Bennett 1971). Positive scaling between genome size and cell size across terrestrial plants has given rise to numerous studies characterizing the many other phenotypic correlates of genome size independent of variation in genome structure, commonly referred to as “nucleotype” effects, although some of these correlations are disputable after accounting for shared phylogenetic history (Bennett 1971; Cavalier-Smith 1978; 1982; Bennett and Leitch 2005). Correlates of genome size encompass an incredible diversity of plant phenotypes, including, for example, the sizes of plant structures, rates of cell division, rates of physiological processes, and tolerances and responses to abiotic conditions (Table 1).

**Table 1.**
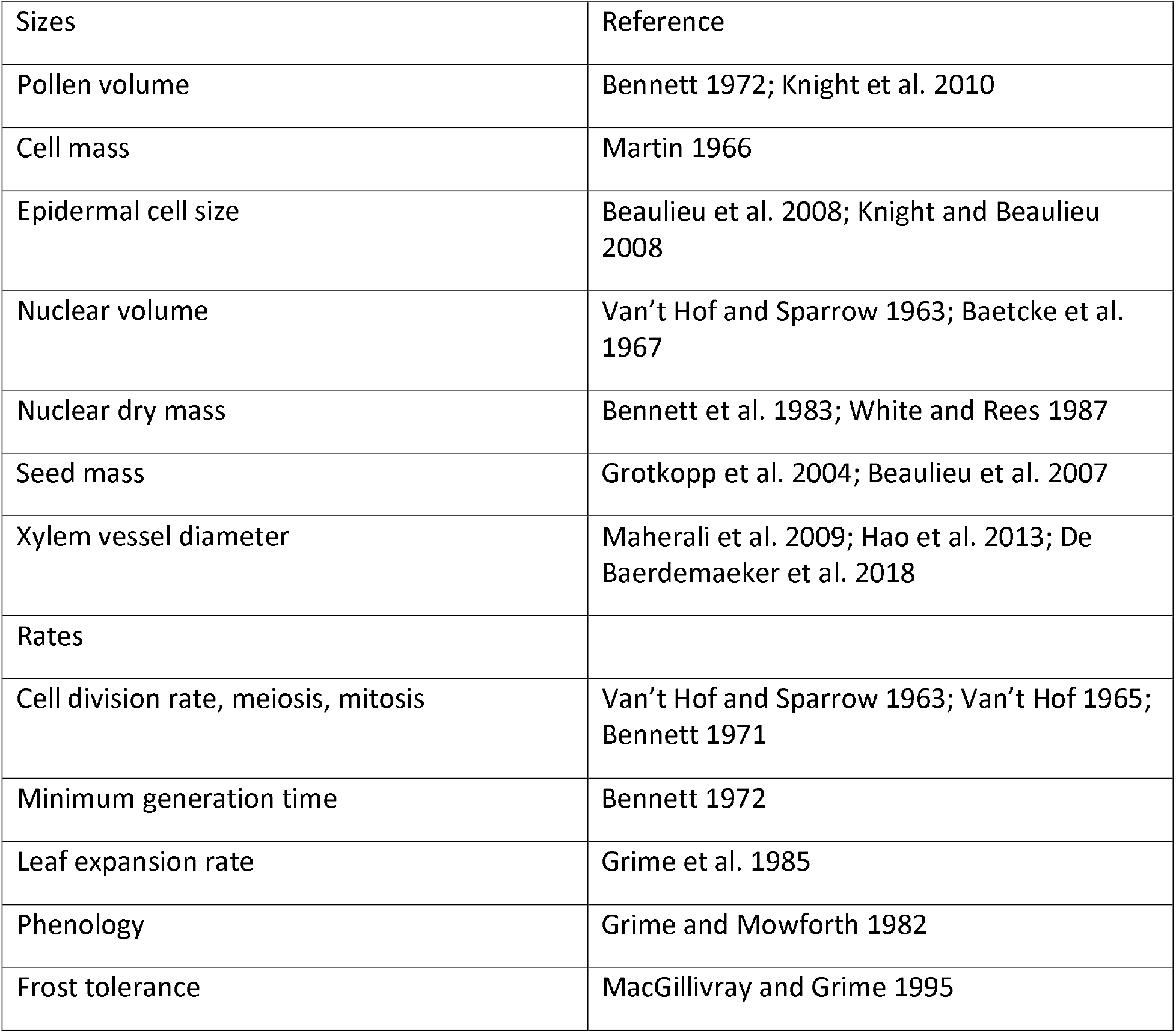
Brief summary of traits shown previously to correlate with genome size.

| Sizes | Reference |
| --- | --- |
| Pollen volume | Bennett 1972; Knight et al. 2010 |
| Cell mass | Martin 1966 |
| Epidermal cell size | Beaulieu et al. 2008; Knight and Beaulieu 2008 |
| Nuclear volume | Van't Hof and Sparrow 1963; Baetcke et al. 1967 |
| Nuclear dry mass | Bennett et al. 1983; White and Rees 1987 |
| Seed mass | Grotkopp et al. 2004; Beaulieu et al. 2007 |
| Xylem vessel diameter | Maherali et al. 2009; Hao et al. 2013; De Baerdemaeker et al. 2018 |
| Rates |  |
| Cell division rate, meiosis, mitosis | Van't Hof and Sparrow 1963; Van't Hof 1965; Bennett 1971 |
| Minimum generation time | Bennett 1972 |
| Leaf expansion rate | Grime et al. 1985 |
| Phenology | Grime and Mowforth 1982 |
| Frost tolerance | MacGillivray and Grime 1995 |

Our goal is not to recapitulate the many reviews about the nucleotype-phenotype relationship but, instead, to align these studies more systematically with the field of plant functional biology. We believe that the diverse impacts of genome-cellular allometry on the body plan of terrestrial vascular plants strongly influences the coordination between plant functional traits and, ultimately, whole organism form-function relationships. Here we summarize previous research, perform new analyses of existing data, and present new data to show how genome size may, through its impacts on cell size and tissue structure, determine the biophysical limits of plant metabolic rates, and, therefore, influence other aspects of ecology and evolution. That genome size may be a key functional trait is not a new idea (Grime 1998). Yet, despite numerous reports of the phenotypic and ecological correlates of genome size (Table 1), it has not been fully integrated into the functional trait literature. Our goal, therefore, is to more directly show how genome size influences plant traits that impact maximum rates of photosynthetic metabolism. Metabolism is central to all three aspects of plant fitness, providing the carbon necessary for allocation to growth, reproduction, and survival. As such, genome size may not itself be a functional trait but instead may define the limits of variation in numerous other functional traits.

## Genome-cellular allometry limits rates of resource transport and metabolism

### Allometry of genome size and cell size

The role of the genome in limiting cell size has been postulated since at least the late 1800s (von Sachs 1893) and was critical in shaping early modern views of the evolution of plant vascular systems (Bailey and Tupper 1918). At a minimum, a cell must contain its genome, and there is a strong relationship between the volumes of meristematic cells and genome size (Šímová and Herben 2012). Cellular expansion from this meristematic minimum size is cell type-specific (Doyle and Coate 2019). Within a cell type, size can be influenced by various environmental and developmental factors (Melaragno et al. 1993). Despite this substantial growth in cell volume during development, there remains a significant effect of genome size on cell size, particularly for stomatal guard cells (Beaulieu et al. 2008; Knight and Beaulieu 2008; Lomax et al. 2013; Simonin and Roddy 2018). For example, stomatal guard cell size and density, which regulate the fluxes of water and CO_2_ between the biosphere and atmosphere, vary within species depending on light, water availability, and atmospheric CO_2_ concentration (Hetherington and Woodward 2003; Franks and Beerling 2009). Furthermore, in the vascular transport network, the sizes of xylem conduits and their density in the leaf are also affected by variation in genome size (Maherali et al. 2009; Hao et al. 2013; De Baerdemaeker et al. 2018; Simonin and Roddy 2018). Yet why genome size and final cell size are correlated within a cell type remains unclear (Doyle and Coate 2019).

We tested whether smaller genomes allow not only for smaller initial and final cell sizes but also for a greater range in final cell size using published data for terrestrial C3 plants. We used data for stomatal guard cells because they are the most commonly measured cell sizes in plants and because their sizes and abundance determine the leaf surface conductance to CO_2_ and water vapor and, therefore, directly control rates of resource transport for use in photosynthetic metabolism. Sizes of guard cells for angiosperms (Beaulieu et al. 2008), gymnosperms, and ferns were compiled previously by Simonin and Roddy (2018), and here we include data for mosses and hornworts from Field et al. (2015) and Renzaglia et al. (2017). We assumed that stomatal guard cells are shaped as capsules, which are composed of a central cylinder with hemispherical ends, such that cell volume could be estimated from cell length as:

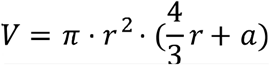

where *r* is the radius of the cylinder and of the hemispherical ends and *a* is the height of the cylinder. We assumed that *a* is equal to 2*r*, such that the guard cell length is equal to 4*r*. Simplifying this equation allowed cell volume to be calculated from guard cell length as:

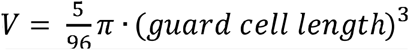

The dumbbell-shaped guard cells present among monocots would likely violate these assumptions about cell shape and so we excluded from this analysis data for the Poaceae, hich are known to have dumbbell-shaped guard cells. Data for meristematic cell volume and genome size were taken from Šímová and Herben (2012). We used linear regression (R package *stats*) to fit the mean response and quantile regression (R package *rq*) to test whether there was greater variation in cell volume among taxa with smaller genomes (i.e. heteroskedasticity), based on differences between quantile regression slopes, using the functions ‘rq’ and ‘anova.rq’.

Across over two orders of magnitude in genome size, meristematic cell volume defined the lower limit of guard cell volume (Figure 1); the smallest guard cells were only slightly larger than meristematic cells of the same genome size. Genome size was a strong and significant predictor of meristematic cell volume (log(volume) = 0.69 ⍰ log(genome size) + 2.68; *R*^*2*^ = 0.98, P < 0.001; Šímová and Herben 2012). Though it explained less of the variation, genome size was a significant predictor of final guard cell volume among terrestrial vascular plants (log(cell volume) = 0.55 ⍰ log(genome size) + 3.44; *R*^*2*^ = 0.48, P < 0.001). Including mosses and hornworts, however, substantially reduced the explanatory power of genome size on cell volume to under 10%. Quantile regression revealed that for vascular plants the slope through the 10th quantile was steeper (slope = 0.66 ± 0.07, intercept = 2.98 ± 0.07) than the slope through the 90th quantile (0.47 ± 0.09), although this difference was not significant (P = 0.07). While there was no significant difference between the 10% and 90% quantile slopes, lower quantiles had consistently steeper slopes when considering all species and also angiosperms alone (Figure S1), suggesting that the smaller minimum cell size allowed by smaller genomes enables greater variation in final cell size. In fact, for a given genome size, interspecific variation in mature guard cell volume could vary by as much as two orders of magnitude among vascular plants. Theoretically, maximum cell size is not as tightly constrained by genome size, such that other cell types can be much larger than guard cells. The greater variation among species with smaller genomes implies that smaller genomes allow for greater plasticity in cell sizes and cell packing densities which directly influence maximum rates of leaf surface conductance to CO_2_ and water and ultimately photosynthetic metabolism per unit leaf surface area (Simonin and Roddy 2018). Further, the greater diversity of cell sizes observed in plants with small genomes suggests that the correlation between genome size and cell size is simply the result of occupying available space within the cell. A small genome can be housed in either a small or a large cell, but a large genome cannot be housed in a cell smaller than its nucleus.

**Figure 1.**
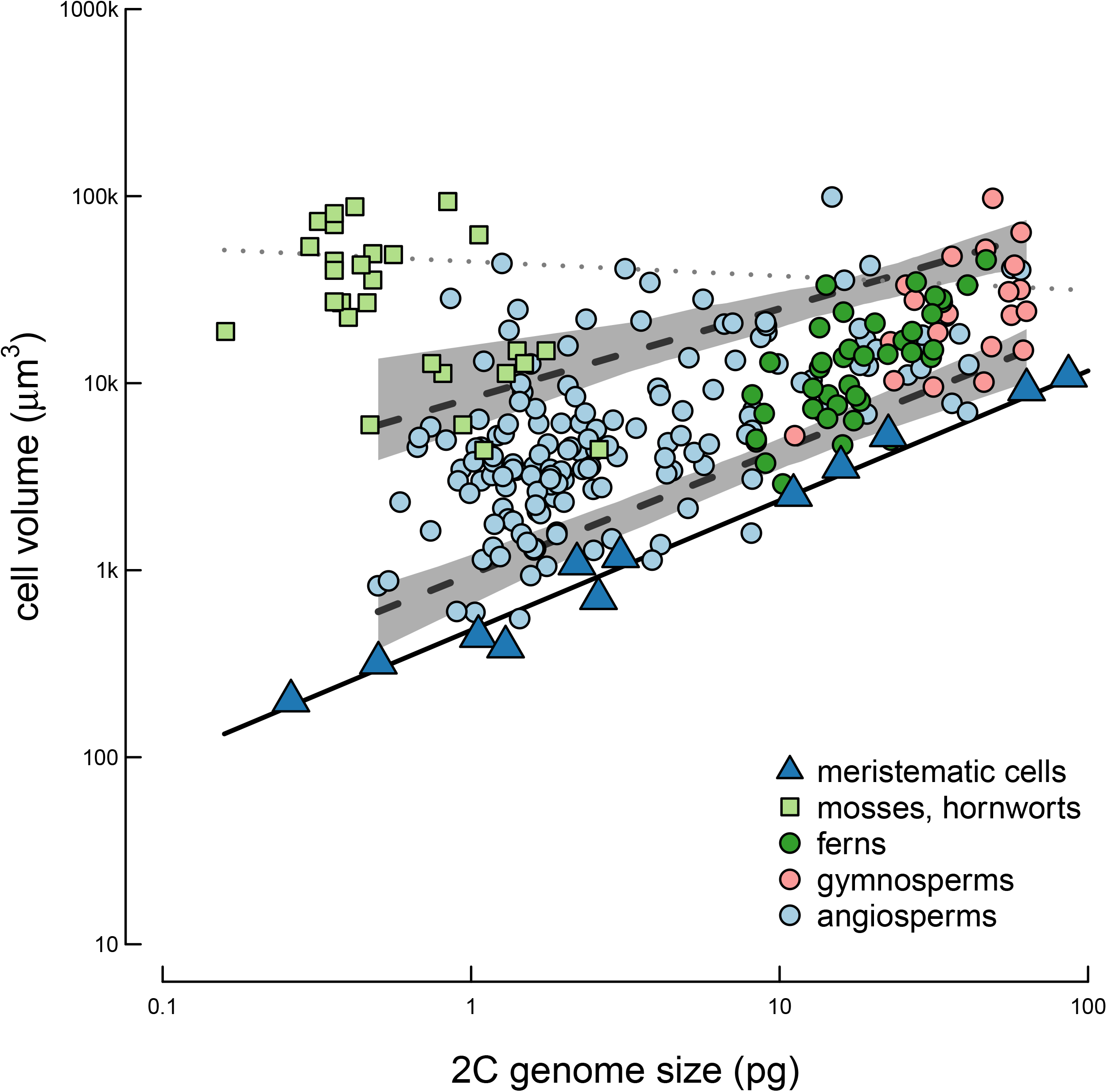
Genome size determines the minimum size of cells, and smaller genomes enable greater variation in final cell size. Data for meristematic cells (blue triangles) were taken from Šímová and Herben (2012), and the solid black line is the regression through these points. Data for mature stomatal guard cells of extant plants (circles and squares) for ferns (dark green), gymnosperms (pink), and angiosperms (light blue) were taken from Simonin and Roddy (2018), and data for mosses and hornworts (light green) were taken from Field et al. (2015) and Renzaglia et al. (2017). The two dashed lines represent the 10th (lower) and 90th (upper) quantile regressions through mature guard cell data for vascular plants with their respective confidence intervals shaded. The dotted line represents the 90^th^ quantile through all guard cell data (vascular and non-vascular plants).

The greater variation in cell volume allowed by smaller genomes (Figure 1) further suggests that smaller genomes allow for greater variation in cell packing densities. For guard cell lengths, stomatal densities, and vein densities, smaller genomes allowed for greater variation in traits across ferns, gymnosperms and angiosperms (Simonin and Roddy 2018). Species with smaller genomes in these datasets are predominantly angiosperms, and these analyses compared distantly related species. We further tested for greater variation in cell sizes and packing densities with smaller genomes among closely related species using taxa in *Rhododendron* (Ericaceae) sect. *Schistanthe* Schltr. (= sect. *Vireya* Blume) and a collection of deciduous *Rhododendron* cultivars that vary in ploidy from diploids to hexaploids. The monophyletic *Schistanthe* clade has a stepwise phylogeographic history, having radiated eastward from the Malay Peninsula and reached New Guinea within the last 15 Ma (Goetsch et al. 2011). We sampled leaves from 19 taxa growing under common garden conditions at the Rhododendron Species Foundation Botanical Garden in Federal Way, WA, USA. Genome sizes were measured following standard protocols (Dolezel et al. 2007) at the Benaroya Research Institute in Seattle, WA, USA. For measurements of stomatal size and density, epidermal impressions were made on fresh leaves using dental putty (Coltene Whaledent President Light Body), transferred using clear nail polish, mounted in water, and imaged using a light microscope.

Measurements of leaf vein density were made on leaf sections cleared by soaking in 4% NaOH, 3% sodium hypochlorite, stained with 1% Safranin O, counterstained with 1% Fast Green, mounted in ethanol, and imaged with a light microscope. Stomatal traits were averaged across ten images per taxon, and leaf vein density was averaged across five images per taxon. Genome sizes for the *Rhododendron* cultivars were measured at the University of Coimbra, Portugal, and all anatomical measurements were made on leaf sections cleared in 4% NaOH, stained in 1% Safranin and mounted in ethanol and Cytoseal (Fisher Scientific). The two datasets of congeners were pooled in statistical analyses. Quantile regression through the 10th and 90th percentile of the species means were used to quantify the variation in traits associated with variation in genome size. Consistent with previous results across terrestrial vascular plants (Simonin and Roddy 2018), among *Rhododendron* taxa, there was greater variation in the sizes and packing densities of veins and stomata among species with smaller genomes (Figure 2). This was apparent due to significant differences between the 10^th^ and 90^th^ quantiles for guard cell length (10^th^: 2.40 ± 1.14, 90^th^: −0.72 ± 1.06; F = 7.11, P < 0.01) and for stomatal density (10^th^: 2.99 ± 10.63, 90^th^: −24.51 ± 12.41; F = 5.90, P = 0.02), but not for vein density (10^th^: 0.14 ± 0.20, 90^th^: −0.36 ± 0.19; F = 3.22, P = 0.07). Further corroborating the significant differences between the 10^th^ and 90^th^ quantile slopes were the more negative slopes among higher quantiles of the data for all traits (Supplementary Figure S2), consistent with the results for guard cell volume among both angiosperms and vascular plants (Figures 1, S1). Thus, across phylogenetic scales, smaller genomes allow for greater variation in the sizes and packing densities of cells.

**Figure 2.**
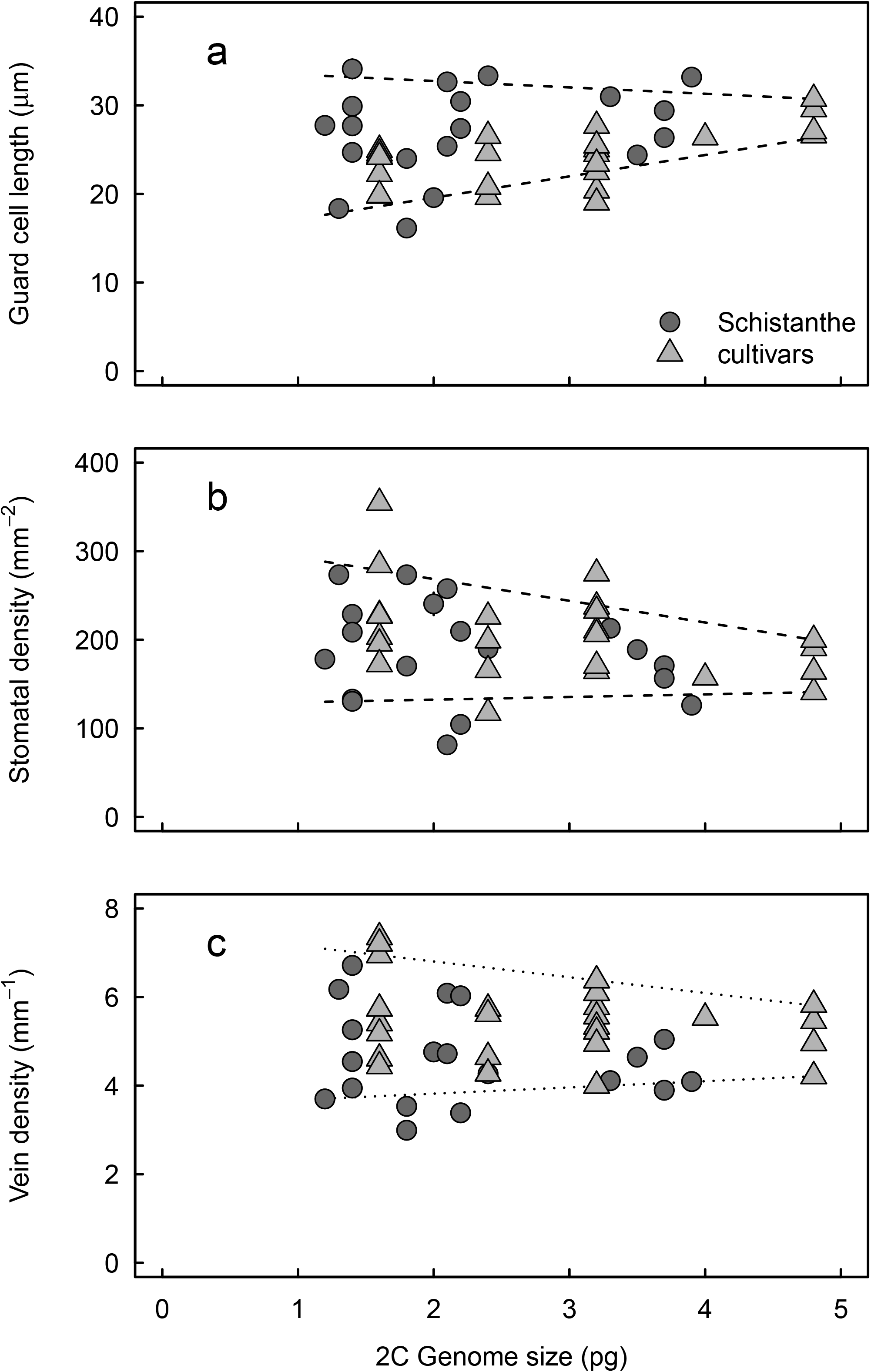
Variation in the sizes and packing densities of stomatal guard cells and leaf veins with variation in genome size among *Rhododendron* sect. *Schistanthe* species (circles) and polyploid *Rhododendron* cultivars (triangles). Lines represent regressions through the 90th (upper) and 10th (lower) quantiles. These quantile regression were significantly different for guard cell length and stomatal density (dashed) but not for vein density (dotted). Genome size limits the lower limit of cell size and the upper limit of cell packing densities, and there is greater variation in anatomical traits among species with smaller genomes.

### Genome size limits maximum photosynthetic metabolism

A major limitation on photosynthetic capacity is the ability to deliver resources to, and export products from, the sites of metabolic processing (Enquist et al. 1998; West et al. 1999a; Brown et al. 2004). At the level of an individual cell–the fundamental unit of living organisms–rates of resource transport are strongly influenced by cell size because the ratio of cell surface area to cell volume increases exponentially with decreasing cell size. Because genome size constrains minimum cell size and the maximum packing densities of cells (Figures 1–2), genome size is predicted to limit the maximum rate of photosynthetic metabolism across vascular plants.

Previous work has hypothesized that genome size would be linked to maximum photosynthetic rate but found little support (Knight et al. 2005; Beaulieu et al. 2007). One major reason for not finding support is that these previous studies attempted to predict variation in *A*_*mass*_, which accounts for the construction costs of leaves, rather than *A*_*area*_, which is the maximum metabolic rate regardless of the construction costs. As described above, *A*_*area*_ would define the maximum amount of carbon assimilated, but how the plant allocates the total assimilated carbon–to growth, reproduction, defense, more durable leaves, etc.–would reflect the numerous factors that influence plant form and other aspects of plant function (Bazzaz et al. 1987). Thus, *A*_*area*_, which is orthogonal to SLA and *A*_*mass*_ (Wright et al. 2004), is predicted to be constrained by cell and genome sizes. Consistent with this prediction, genome size is a strong predictor of the sizes and densities of stomatal guard cells and leaf veins across vascular plants (Simonin and Roddy 2018), and we predicted, therefore, that genome size would, via its effects on the sizes and packing densities of cells, limit *A*_*area*_. It is important to clarify that many factors can influence *A*_*area*_ of a given leaf. For example, nutrient deficiency and water stress can reduce *A*_*area*_ below its theoretical maximum–independent of the effects of cell and genome size–by limiting either the biochemical or stomatal contributions to carbon assimilation. When these other factors are not limiting, then cell size is predicted to limit *A*_*area*_, and, as a result, we predicted that genome size would define the upper limit (estimated using quantile regression) of *A*_*area*_.

Data for area-based maximum photosynthetic rate were compiled from the primary literature (Supplemental Table 1) and merged with the Kew Plant DNA C-Values Database (Bennett and Leitch 2012). This dataset included 210 species, of which 138 were angiosperms, 46 were gymnosperms, and 26 were ferns. We tested whether genome size limits *A*_*area*_ using quantile regression. Like above, we estimated the upper limit of *A*_*area*_ as the 90^th^ quantile, but include slope estimates across quantiles (Figure S3). Standard errors around these quantile slopes were estimated by bootstrapping 300 replicates. There is no phylogenetically corrected method for estimating quantile slopes, so we tested whether the pattern observed across all species was also apparent only among the angiosperms, which exhibit the largest range in genome size of the three main groups of vascular plants. This analysis helped to determine whether the effects of genome size on *A*_*area*_ were driven solely by the divergences between the three major clades.

Smaller genomes enabled higher maximum photosynthetic rates across and within major plant clades (Figure 3). Across all terrestrial vascular plants, the upper limit (the 90th quantile) of *A*_*area*_ was defined by genome size (slope = −0.18 ± 0.03). A nearly identical slope of the 90th quantile was apparent only among the angiosperms (−0.19 ± 0.05), suggesting that the effect of genome size on maximum *A*_*area*_ was not due solely to the divergences between the three major clades. Across all quantiles there was little difference between the quantile slopes estimated for all species versus the angiosperms alone, and these quantile slopes were mostly within the confidence interval of the regression slope through the entire dataset (Figure S3).

**Figure 3.**
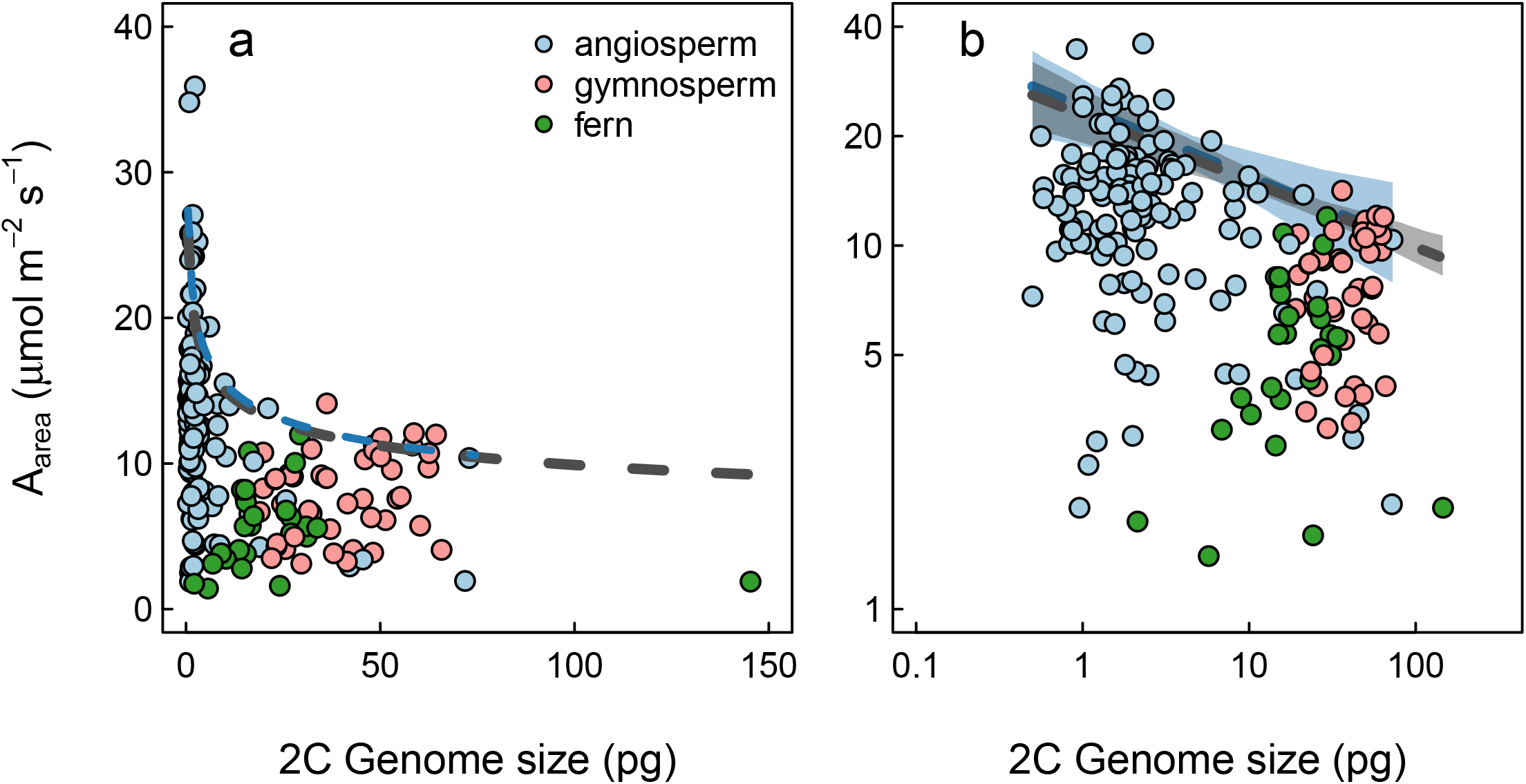
Genome size limits the maximum rate of photosynthesis (*A*_*area*_) across C3 terrestrial plants. (a) Untransformed relationship and (b) log-transformed relationship. Dashed black lines are regressions through the upper 90th quantile of all data with grey shading representing the 95% confidence interval. Blue dashed lines and blue shading represent the 90^th^ quantile regression and its 95% confidence interval for angiosperms alone, showing that the same slope defines the upper limit among only the angiosperms as across all three major clades of vascular plants.

The scaling relationship between *A*_*area*_ and genome size follows naturally from the relationships between genome size and the sizes and densities of veins and stomata. However, veins and stomata are not the only cells responsible for driving variation in photosynthetic rates. While the maximum rate of CO_2_ diffusion into the leaf is defined by the sizes and densities of stomata (Franks and Beerling 2009), once inside the leaf, CO_2_ must diffuse through the leaf intercellular airspace and into the chloroplasts lining the interior surfaces of mesophyll cells. Thus, the three-dimensional structure and organization of the mesophyll is predicted to be a prime target for selection on photosynthetic metabolism (Tholen et al. 2012; Ren et al. 2019) and to be critical to leaf photosynthetic function (Earles et al. 2019). The limited evidence on *Arabidopsis thaliana* mutants suggests that cell size is critical to this mesophyll architecture (Lehmeier et al. 2017). Based on the results presented here (Figure 3) and elsewhere (Simonin and Roddy 2018), we predict that the scaling relationships between genome size and cell size that coordinate veins and stomata extend also to the sizes of cells and their organization within the leaf mesophyll.

### Genome size may limit the rate of metabolic up- or down-regulation

Although maximum potential rate of leaf gas exchange is an important parameter determining a species’ physiological capacity, the actual rate of leaf gas exchange at any given moment is often substantially lower, depending on a variety of physiological and environmental factors (e.g. light level, atmospheric humidity, leaf temperature, plant water status). Changes in sun angle, shading by passing clouds, and self-shading by fluttering leaves all drive changes in incoming solar radiation, and these rapid dynamics have influenced the evolution of photosynthetic biochemistry (Pearcy 1990). Under naturally varying conditions, leaf gas exchange fluctuates dramatically and rarely reaches its maximum rate, with greater variation occurring at the top of the plant canopy. How frequently a leaf can reach its maximum gas exchange rate and how well it can optimize its physiological processes to environmental conditions depend on how rapidly the leaf can respond to dynamic, fluctuating conditions.

There is an emerging consensus that smaller stomata respond more rapidly to fluctuating conditions than larger stomata, allowing leaves with smaller stomata to more closely tune their physiological rates with environmental conditions (Drake et al. 2013; Lawson and Blatt 2014; Lawson and Vialet-Chabrand 2019). Leaf physiological processes change at different rates, with changes in stomatal conductance occurring an order of magnitude more slowly than changes in photosynthesis (McAusland et al. 2016). This difference in response times between physiological processes (e.g. photosynthetic assimilation rate and stomatal conductance) can reduce water use efficiency when stomata are closing and reduce photosynthetic efficiency when stomata are opening (Lawson and Vialet-Chabrand 2019), limiting total photosynthesis by up to 20% (Lawson and Blatt 2014). If stomatal response times are directly limited by the size of stomata then genome-cellular allometry may limit not only the maximum rate of metabolism but also how quickly metabolism can respond to fluctuating environmental conditions. Of the species for which stomatal response times were measured by McAusland et al. (2016) and Drake et al. (2013), twelve were included in the Kew Plant DNA C-Values database. Consistent with previous reports, there was a positive correlation between genome size and guard cell length (*R*^*2*^ = 0.36, P < 0.05; Figure 4a), and stomatal response rate exhibited a triangular relationship with genome size such that smaller genomes exhibited both higher maximum stomatal response rates but also a greater variation in stomatal response rate. While the available data on stomatal response rates measured using standard protocols are limited, these preliminary results suggest that genome size indirectly limits the maximum rate of stomatal opening and closing via its effects on the sizes and densities of stomata.

**Figure 4.**
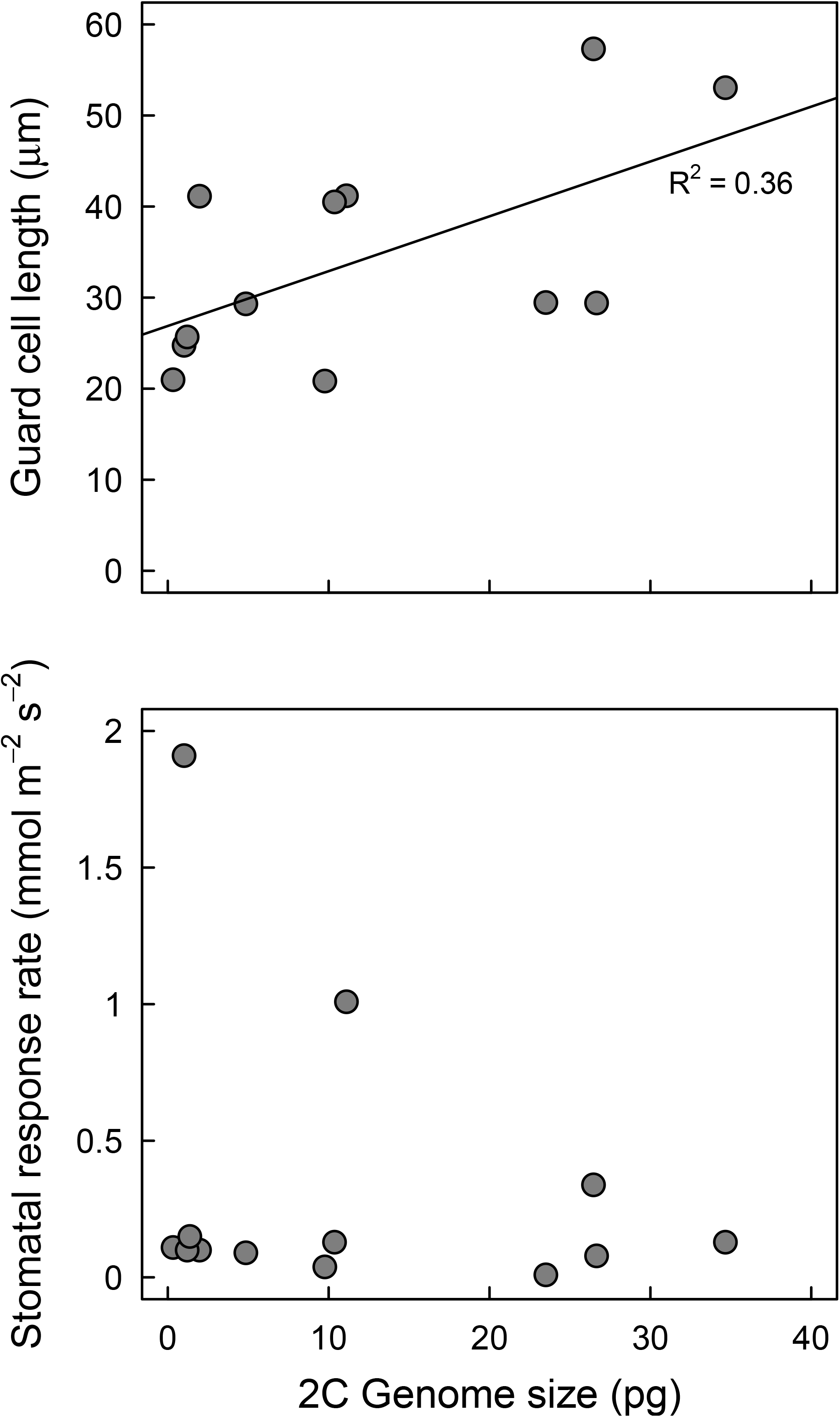
Genome size may limit the maximum rate of stomatal response (i.e. how fast stomata can open or close). Data taken from McAusland et al. (2016) and Kew Plant DNA C-values Database.

## How genome size-metabolism scaling may impact plant biogeography

### Polyploidy thought to increase niche breadth

Variation in genome size and structure associated with polyploidization has long been considered to be an important driver of plant evolution and to be associated with shifts in environmental tolerances, habitat breadth, trait variation, and interspecific interactions (Stebbins 1940; Otto and Whitton 2000; Soltis et al. 2003; Soltis et al. 2014; Barker et al. 2016a,b), and niche differentiation between polyploids and their diploid parentals has been considered a prerequisite for the successful establishment of newly arisen polyploids (Levin 1975; Fowler & Levin, 1984). Describing the types of polyploids and how they are has been thoroughly reviewed elsewhere (e.g. Stebbins 1947; Soltis et al. 2015), and we focus our discussion here on how and why ploidy–via its relationship with genome size– may or may not correlate with species distributions and habitat breadth. Until they can be more rigorously tested, these ideas will remain speculative.

Polyploids have been hypothesized to be better adapted to extreme habitats, to have greater hardiness, and to have greater ecological adaptability (reviewed by Stebbins 1985: Brochmann et al. 2004). The possible mechanisms for these effects can be roughly grouped into two categories: one involving the genetic and genic content of the polyploid genome and one involving the nucleotypic effects of ploidy and genome size. Because polyploid genomes commonly have additional genome copies, they have higher absolute genic contents, would enable neofunctionalization of duplicated genes, and typically have higher heterozygosity, all of which can promote higher tolerances of environmental conditions. The nucleotypic effects of ploidal variation, though long recognized (Stebbins 1940), are often confounded with nucleotypic effects of genome size variation.

While ploidy and genome size are commonly assumed to be synonymous, at broad phylogenetic scales there is generally no relationship between genome size and ploidy (Leitch and Bennett 2004), reflecting the complex history of both ancient and contemporary whole genome duplications, particularly among the angiosperms (Jiao et al. 2011; Clark and Donoghue 2018; Landis et al. 2018; Ren et al. 2018). In contrast to pteridophytes, which also frequently undergo whole genome duplications (Clark et al. 2016), angiosperm genomes readily rediploidize after polyploidization such that genome size and ploidy are positively correlated only for narrowly defined phylogenetic groups (i.e. within genera and families, Figure 5; Leitch and Bennett 2004; Dodsworth et al. 2016). If leaf and plant structure and function influence ecological tolerances and habitat breadth (i.e. if plant structure-function is adaptive), then the nucleotypic effects of genome size are predicted to influence environmental tolerances.

**Figure 5.**
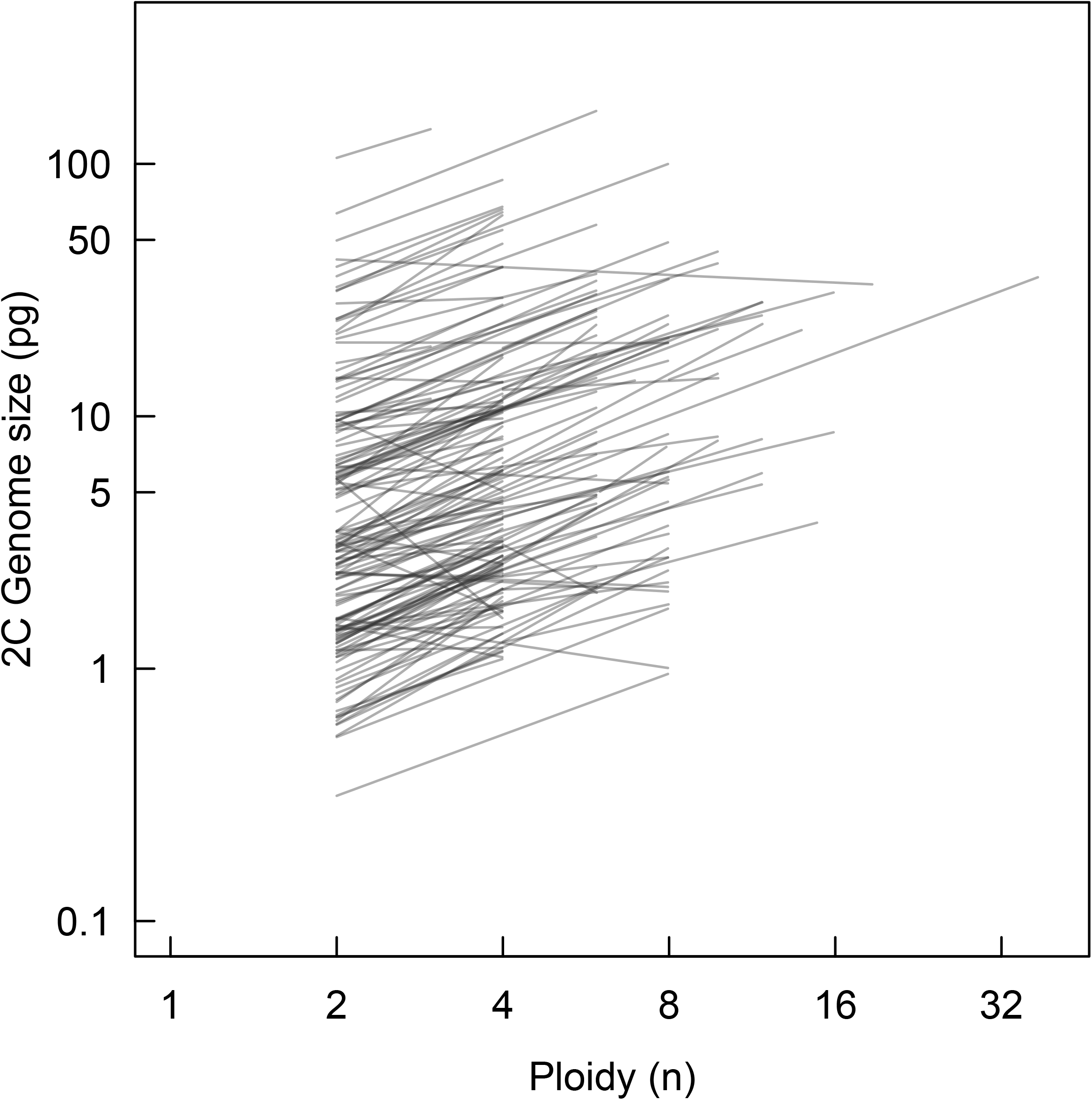
Relationship between genome size and ploidy for angiosperms. Each line represents the linear regression within a genus. At narrow taxonomic scales, ploidy and genome size are correlated, but at broad taxonomic scales (i.e. among all angiosperms), there is no relationship between genome size and ploidy due to rediploidization.

### Smaller genomes enable greater phenotypic plasticity

One long-standing hypothesis is that higher ploidy is related to wider habitat breadth because polyploids can tolerate greater ecological stress. Higher ploidy is associated with greater heterozygosity (i.e. greater genetic diversity) and, frequently, higher genic content due to multiple genome copies, both of which are thought to promote plasticity and enable polyploids to withstand a greater range of environmental conditions than diploids. However, several studies testing this hypothesis have not observed polyploids to have greater habitat breadth (e.g. Stebbins 1985; Martin and Husband 2009; Glennon et al. 2014; Johnson et al. 2014). Furthermore, these tests frequently find that diploids exhibit greater habitat breadth than polyploids (Petit and Thompson 1999; Hijmans et al. 2007; Brittingham et al. 2018; Castro et al. 2019). One reason is that traits are not necessarily more variable in polyploids than in diploids (Stebbins 1985; Wei et al. 2018).

We predict that one reason ploidy is not commonly found to correlate with ecological breadth is because genome size–rather than ploidy per se–drives variation in the absolute range of potential cell sizes and, by extension, phenotypic plasticity in rates of resource transport and metabolism. Thus, the phylogenetic scale-dependence of the relationship between genome size and ploidy (Figure 5), particularly among the angiosperms, could lead to confounding patterns depending on the phylogenetic scale at which comparisons are made. For example, in the analysis of Rice et al. (2019), ploidy was determined relative to other closely related species, such that within genera or families ploidy and genome size are positively correlated, suggesting that the bias towards higher abundances of polyploids at higher latitudes may reflect nucleotypic effects of genome size on cell size and metabolism. The complex, fluctuating process of polyploidization and rediploidization, which can winnow the genome nonrandomly (Wendel 2015), would promote the proliferation of beneficial elements associated with genome duplications (e.g. more gene copies that can neofunctionalize) while reducing the size of the genome needed to maintain high rates of development and metabolism (Table 1).

We posit here that the nucleotypic effects of genome size, regardless of ploidy, may influence environmental tolerances. Because smaller genomes allow for greater variation in cell size and metabolism (Figures 1–3), species with smaller genomes may be better able to fine tune their tissue structure to environmental conditions. This flexibility would allow species with smaller genomes to better optimize their metabolic rates in order to occupy a wider range of environmental conditions. Combined with the effects of genome size on rates of cell division (Van’t Hof and Sparrow 1963; Van’t Hof 1965; Šímová and Herben 2012), the greater plasticity in cell size and higher metabolic rates attainable by species with small genomes may enable them to better colonize new habitats.

### Community-scale patterns in genome size across gradients in productivity

If habitats filter species based on rates of metabolism and if there are nucleotypic effects of genome size on metabolism, then community-scale distributions of genome size may vary across gradients of productivity. In habitats that can support high rates of productivity and primary metabolism, species with small genomes are expected to predominate because they can maintain higher rates of metabolism and more rapidly adjust their physiology to match environmental conditions. This strategy would be one of maintaining steady state physiological processes. At a broad scale, this prediction holds because angiosperms, which have, on average, smaller genomes than other vascular plants are dominant in most ecosystems, particularly those characterized by high productivity. However, high rates of metabolism and maintaining steady-state physiology, even among the angiosperms, are not always favorable. Two such habitats are those characterized by extreme water and nutrient limitation, such as deserts and epiphytic habitats, and by extreme cold, such as high latitudes. Higher incidences of polyploids have been commonly reported in higher latitudes and among arctic floras (Brochmann et al. 2004; Rice et al. 2019), but arid habitats have received less attention.

Arid and epiphytic habitats are characterized by low productivity and may support species with large genomes. In these habitats, high rates of metabolism are not always favored, which may relax selection for small genomes. One strategy common in arid and epiphytic habitats is succulence, which is often associated with Crassulacean acid metabolism (CAM) photosynthesis. The CAM syndrome limits water loss by restricting CO_2_ uptake and water loss to nighttime when humidity is high and the atmospheric demand for evaporation relatively low. As a result, CAM species typically rely more heavily on resource storage (e.g. CO_2_, H_2_O) or non-steady-state physiology to maintain photosynthetic metabolism and limit water loss. If metabolism is one agent of selection on genome size, then we would predict that in arid, resource poor environments, selection for small genomes (associated with small cells and high metabolic rates) may be weak among CAM species, allowing genomes of CAM species to expand in size. We tested this hypothesis using the taxonomic distributions of CAM photosynthesis from Smith and Winter (1996) and genome size data from the Kew Plant DNA C-Values Database (Bennett and Leitch 2012). For C3, we used the broad distribution of angiosperms reported in Simonin and Roddy (2018), which are representative of extant angiosperm diversity. We scored as CAM the narrowest taxonomic level in the Kew DNA C-Values Database that was listed as containing CAM by Smith and Winter (1996). For example, if a genus were listed as containing any CAM species, all species in the genus were assumed to exhibit CAM photosynthesis. This approach was biased against observing differences in genome size between C3 and CAM species because it necessarily grouped some C3 species as CAM. To account for phylogenetic history, we constructed a dated, family-level supertree using the methods described in Simonin and Roddy (2018), and compared C3 and CAM genome sizes using the *phylANOVA* function in ‘phytools’ (Revell 2012). Log-normalized genome sizes were significantly larger among CAM species than among C3 species (t = 8.11, df = 284.03, P < 0.001) even after accounting for shared phylogenetic history (t = 7.51, P < 0.05; Figure 6), consistent with the prediction that large genomes may evolve when selection for high rates of metabolism is weak. However, future analyses that incorporate better determination of the phylogenetic distributions of photosynthetic pathways is needed to more rigorously test whether the evolution of CAM photosynthesis and its associated switch towards non-steady-state physiological processes is indeed associated with increases in genome size.

**Figure 6.**
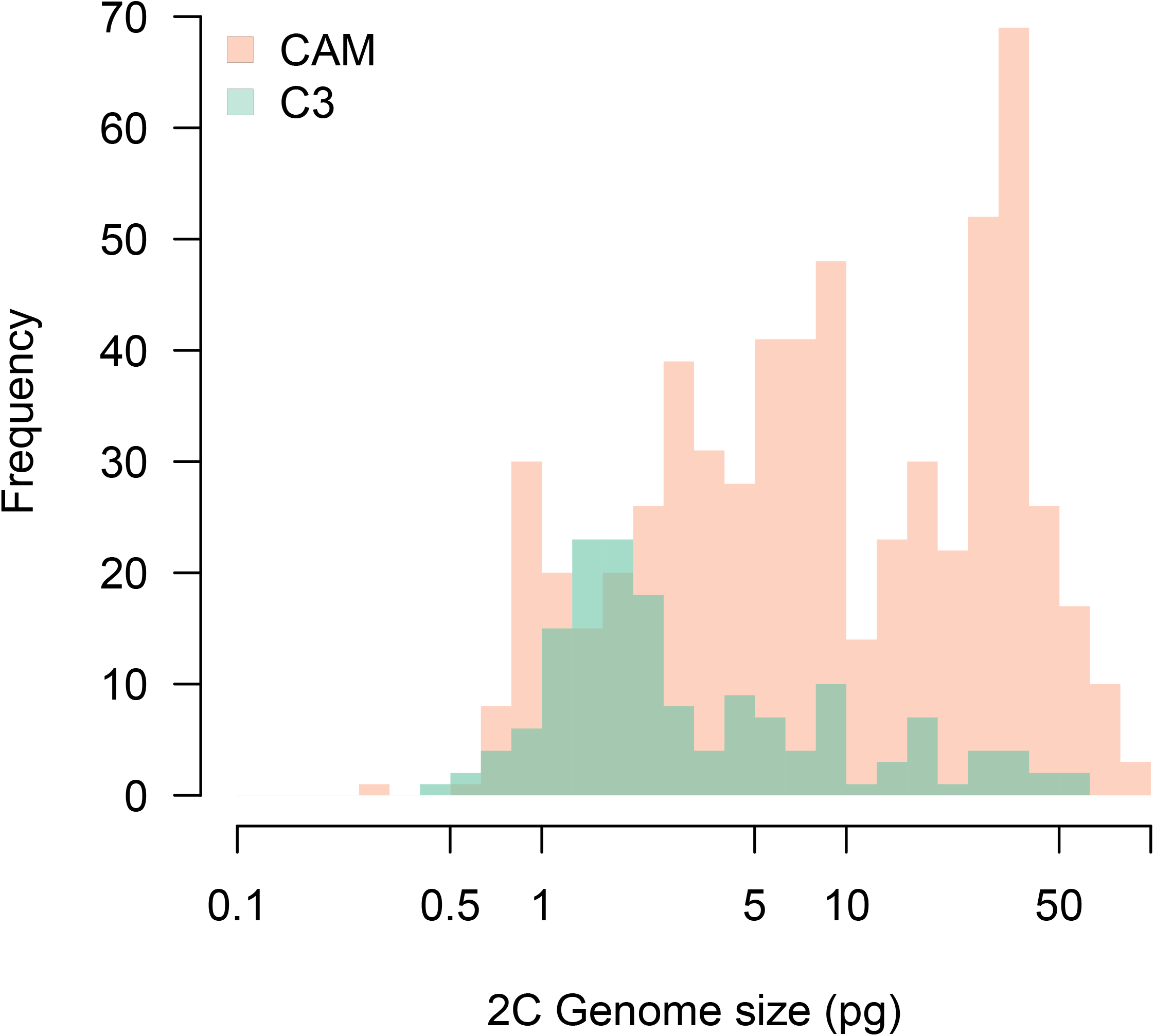
Distributions of genome size for C3 and CAM species show CAM lineages have significantly larger genomes than C3 lineages. Lineages identified as CAM likely include many C3 species; see text for details on identification of photosynthetic pathways. There was a significant difference in log-normalized genome size for the two photosynthetic pathways, even after accounting for shared phylogenetic history.

Arid, resource poor habitats are not exclusively composed of species with large genomes. Rather, they may harbor a diversity of strategies associated with divergent niches. In deserts, physiological strategies can be arrayed along a spectrum from strict non-steady-state physiology characterized by low rates of metabolism (e.g. obligate CAM) to quasi-steady-state physiology (e.g. C3 species) characterized by high rates of metabolism (Nobel and Jordan 1983; Hunt and Nobel 1987). While CAM species can rely on resource storage during periods of limited water availability, C3 species in deserts tend to function during a relatively narrow period of time when water is available. Thus, because their carbon gain is limited to such a short time period, C3 desert plants may have small genomes and cells that enable high rates of metabolism. In fact, desert shrubs have the highest rates of stem hydraulic conductance measured in C3 plants (Mencuccini 2003), and even among species from humid tropical forests, dry forest species have higher hydraulic conductance than wet forest species (Brenes-Arguedas et al. 2013). Thus, less productive habitats may select not simply for larger genomes but instead allow for multiple strategies that encompass a broader range of metabolic rates and, by extension, greater variation in genome size at the community level.

### Smaller genomes increase the probability of invasiveness

The multifaceted effects of genome size on plant structure, function, and ecology (Table 1) is particularly relevant to the study of invasive species. Identifying the traits that allow an introduced species to establish, naturalize, and invade into a new environment is a central aim of invasion biology (Simberloff 2011), with broader implications for plant biogeographic patterns. Here we distinguish between nonnative species–those that survive and reproduce in their introduced range–and nonnative invasive species–those that can disperse, establish, and spread far from their original source of introduction (Richardson et al. 2011). This distinction is important because prior studies on the traits of ‘invaders’ focus on these different subsets of species, which have slightly different, but overlapping, sets of traits that determine whether they can survive and reproduce versus invade non-native regions (Kleunen et al. 2015).

Early theory on the distinguishing traits of invasive plants postulated that “ideal weeds” should grow rapidly, produce seed continuously and in high number throughout the growing season, be tolerant to a wide range of environmental conditions, exhibit high trait plasticity, and be able to reproduce vegetatively from fragments (Baker 1974). On average, these predictions have been upheld, with nonnative invasive plants tending to exhibit traits consistent with high fitness (e.g. number of flowers, fruits, or seed or germination rates), high relative growth rates, high dispersal abilities (e.g. smaller seeds), and more efficient carbon-capture strategies (e.g. high specific leaf area), relative to co-occurring native species (Leishman et al. 2007; Kleunen et al. 2010; Ordonez et al. 2010; Kuester et al. 2014) or naturalized but not invasive nonnative species (Rejmánek and Richardson 1996; Gallagher et al. 2014). Combined, these traits confer a growth advantage, such that plants with small seeds can disperse further distances, have shorter generation times, and higher relative growth rates, owing to the greater rates of cell division and higher metabolic rates provided by smaller genomes (Pandit et al. 2014; Suda et al. 2015). Indeed, even within species, populations with smaller genomes are more likely to successfully invade new habitats (Pysek et al. 2018).

Because many of the traits linked with invasiveness can be influenced by both ploidy and genome size, both have been implicated as underlying features driving invasion (Pandit et al. 2014; Suda et al. 2015). Because polyploids are thought to be better able to tolerate environmental fluctuations and to be better able to adapt to new environments, polyploids tend to be overrepresented among nonnative invasives compared to native angiosperms (Rejmánek and Richardson 1996; Prentis et al. 2008; Beest et al. 2011; Pandit et al. 2014). Similarly, nonnative invasive species tend to have smaller genomes than non-invasive plants (both native and non-native), which is thought to be due to the diverse effects of genome size on metabolism, rates of development and growth, and seed size (Rejmánek and Richardson 1996; Bennett et al. 1998; Kubešová et al. 2010; Pandit et al. 2014). However, the complex, scale-dependent relationship between ploidy and genome size (Figure 5) complicates a clear understanding of the effects of ploidy versus genome size on invasiveness (Rejmánek and Richardson 1996; Pandit et al. 2014). Because angiosperms, which predominate among nonnative invasives, readily rediploidize and downsize their genomes subsequent to whole genome duplications (Leitch and Bennett 2004), assessing the relative effects of ploidy versus genome size on invasiveness can be difficult. For example, the likelihood of being invasive increases with chromosome number and ploidy but decreases with genome size (Rejmánek and Richardson 1996; Pandit et al. 2014). The multiple paths to polyploidization and the selective retention of only certain parts of the genome during subsequent genome downsizing (Wendel 2015) could explain how both higher ploidy and smaller genomes are correlated with invasiveness.

## A possible role for metabolism in genome size evolution

As the major source of energy and matter for the biosphere, photosynthetic metabolism represents a first-order control over ecological processes globally. This fundamental link between metabolic and ecological processes has driven the development of the Metabolic Theory of Ecology (MTE) that provides a mechanistic framework for predicting variation in organismal life history attributes, population dynamics, and larger scale ecosystem processes from organismal-level traits related to resource supply for metabolism (West et al. 1997; Enquist et al. 1998; West et al. 1999a; West et al. 1999b; West et al. 2002; Price et al. 2010). While appealing and seemingly endowed with incredible explanatory power, a number of criticisms of the theory and its assumptions have been consistently raised (Kozłowski and Konarzewski 2004; Kozłowski and Konarzewski 2005; Price et al. 2012). One primary assumption is that the sizes of terminal units in vascular networks (e.g. capillaries in circulatory systems or terminal veins in plant leaves) are invariant. The problems with this assumption have been thoroughly detailed for animal circulatory systems with the allometry of genome size and cell size emerging as a critical factor influencing how body size scales with metabolism (Kozłowski et al. 2003). Furthermore, the allometry of genome size and cell size (Figure 1) and the effects of genome size on maximum metabolic rate (Figure 3) presented here suggest that this assumption is violated in plants, as well. Modifications to the original model that relax some of its assumptions have improved model predictions for plants, particularly by allowing for variation in the packing of xylem conduits (Savage et al. 2010). However, the nucleotypic effects of genome size have yet to be incorporated, although they may further improve models and help to clarify the constraints and major innovations driving botanical form, function, and diversity.

The effects of genome size on cell sizes and packing densities across vascular plants (Figures 1,2; Beaulieu et al. 2008; Simonin and Roddy 2018) and the importance of cell size in metabolism (Savage et al. 2010) together suggest that there may be a role for metabolism in the evolution of genome size. While it is appealing to expect that genome size may predict metabolic rate, the effects of genome size are likely more nuanced. Because genome size defines only the lower limit of cell size, genome size may limit only the maximum possible rate of energy and matter exchange (Figure 3), rather than being a clear predictor of metabolism more generally. This suggests that evolutionary increases in metabolic capacity may be tied to the evolution of genome size, such as has been described in birds (Wright et al. 2014). How selection on genome size *per se* may be translated into alterations of genome sequence structure is unclear but would be an important step towards understanding the drivers of genome size variation. Independent evidence for the role of metabolism in shaping genome-cellular allometry can be evaluated by comparing structures with similar developmental origins such as flowers and leaves (Olson and Pittermann 2019). Flowers, unlike leaves, need not support high rates of energy and matter exchange for use in photosynthetic metabolism and generally have larger cells and lower cell packing densities than their conspecific leaf counterparts (Roddy et al. 2013, 2019; Zhang et al. 2018; Roddy *in press*). Thus, under different selection regimes due to differences in metabolism, traits can diverge even within the same organism (Olson and Arroyo-Santos 2015). Furthermore, defining the biophysical limits of phenotypic variation is central to understanding the diversity of plant form and function, and our analyses suggest that genome size defines one bound to the range of possible cell sizes.

---

Figure S1. Quantile regression slopes and bootstrapped standard errors for of cell volume and genome size data plotted in Figure 1 for vascular plants. Quantiles were calculated for every 5% of the data (5% to 95%) for all vascular plants (ferns, gymnosperms, angiosperms; black points) and for angiosperms only (blue points). Points are jittered horizontally so they do not plot on top of each other. The OLS slope through the entire dataset (solid red line) and its confidence interval (dotted red lines) are included for comparison. Lower quantiles of the data have consistently steeper slopes.

Figure S2. Quantile regression slopes and bootstrapped confidence intervals for (a) guard cell length, (b) stomatal density, and (c) vein density of *Rhododendron* subsect. *Schistanthe* species and *Rhododendron* cultivars. Original data plotted in Figure 2. Quantiles were calculated for every 5% of the data (5% to 95%), with standard errors estimated by bootstrapping 300 replicated. The OLS slope (solid red line) and its confidence interval (dotted red lines) are included for comparison. For all three traits, lower quantiles of the data have consistently steeper slopes.

Figure S3. Quantile regression slopes and bootstrapped standard errors for *A*_*area*_ and genome size data plotted in Figure 3. Quantiles were calculated for every 5% of the data (5% to 95%) for all vascular plants (ferns, gymnosperms, angiosperms; black points) and for angiosperms only (blue points). Points are jittered horizontally so they do not plot on top of each other. The OLS slope through the entire dataset (solid red line) and its confidence interval (dotted red lines) are included for comparison. Lower quantiles of the data have consistently steeper slopes.

