## Supplementary figures and images for "The scaling of genome size and cell size limits maximum rates of photosynthesis with implications for ecological strategies"

### Figure S1

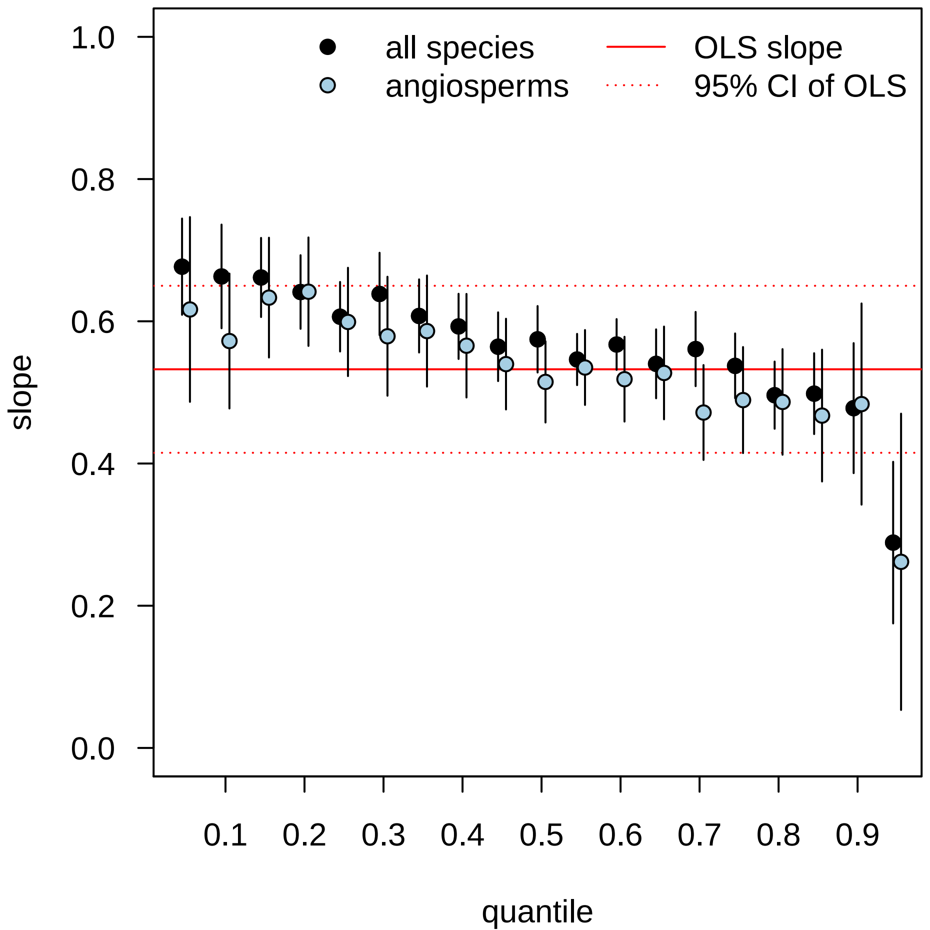

### Figure S2

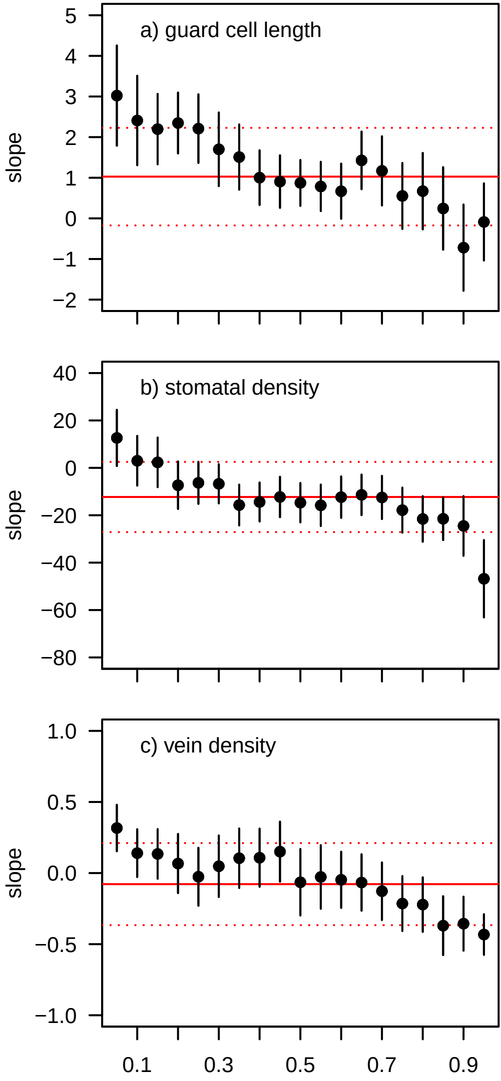

### Figure S3

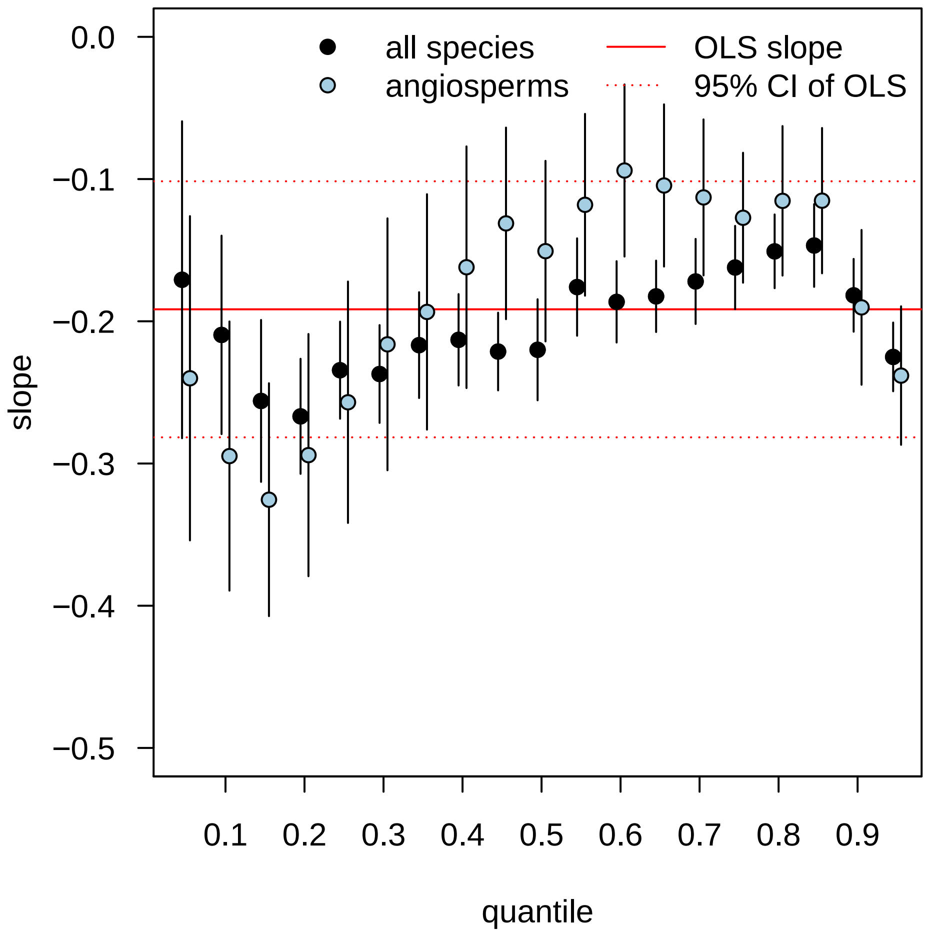
